# Long-term incubation of lake water enables genomic sampling of consortia involving Planctomycetes and Candidate Phyla Radiation bacteria

**DOI:** 10.1101/2021.09.01.458585

**Authors:** Alexander L. Jaffe, Maxime Fuster, Marie C. Schoelmerich, Lin-Xing Chen, Jonathan Colombet, Hermine Billard, Télesphore Sime-Ngando, Jillian F. Banfield

**Affiliations:** Department of Plant and Microbial Biology, University of California, Berkeley, CA; Laboratoire Microorganismes: Génome et Environnement (LMGE), UMR CNRS 6023, Université Clermont-Auvergne, F-63000 Clermont-Ferrand, France; Innovative Genomics Institute, University of California, Berkeley, CA; Department of Earth and Planetary Science, University of California, Berkeley, CA; Department of Environmental Science, Policy, and Management, University of California, Berkeley, CA; Chan Zuckerberg Biohub, San Francisco, CA

## Abstract

Microbial communities in lakes can profoundly impact biogeochemical processes through their individual activities and collective interactions. However, the complexity of these communities poses challenges, particularly for studying rare members. Laboratory enrichments can select for subsystems of interacting organisms and enable genome recovery for enriched populations. Here, a reactor inoculated with water from Lake Fargette, France, and maintained under dark conditions at 4°C for 31 months enriched for diverse Planctomycetes and Candidate Phyla Radiation (CPR) bacteria. We reconstructed draft genomes and predicted metabolic traits for 12 diverse Planctomycetes and 9 CPR bacteria, some of which are likely representatives of undescribed families or genera. One CPR genome representing the little-studied lineage Peribacter (1.239 Mbp) was curated to completion, and unexpectedly, encodes the full gluconeogenesis pathway. Metatranscriptomic data indicate that some Planctomycetes and CPR bacteria were active under the culture conditions. We also reconstructed genomes and obtained transmission electron microscope images for numerous phages, including one with a >300 kbp genome and several predicted to infect Planctomycetes. Together, our analyses suggest that freshwater Planctomycetes may act as hubs for interaction networks that include symbiotic CPR bacteria and phages.

## INTRODUCTION

Freshwater lakes host diverse microbial communities that likely control ecosystem biogeochemistry [1, 2]. Here, we established a laboratory culture based on an inoculum from Lake Fargette, France, and probed enriched populations using a combination of metagenomics, metatranscriptomics, and microscopy. This site was originally chosen for a parallel study of enigmatic ‘aster-like nanoparticles’ (ALNs) [3, 4]. We recovered draft genomes for abundant and transcriptionally active Planctomycetes, as well as CPR bacteria, phages, and eukaryotic viruses. Overall, we provide clues to interactions among microbial groups in a lake ecosystem that were linked by co-enrichment.

## RESULTS

### Community composition and genome reconstruction

Concentrate from the 0.2 μm–25 μm size fraction of the highly eutrophic Lake Fargette, France, was incubated at 4°C in the dark with filtered and sterilized lake water (<20 KDa). After 31 months, DNA and RNA were extracted for metagenomic and transcriptomic analyses. DNA reads were assembled and the contigs profiled to identify ribosomal protein S3 (rpS3). Profiling using the predicted rpS3 protein sequences revealed that the enrichment was bacterially-dominated, although several members of the Thaumarchaeota were present (Fig. 1a, Table S1). The most abundant organisms overall were Planctomycetes (~27% of overall rpS3 coverage), including the most abundant singular organism (Fig. 1, Table S1). CPR bacteria were 5 of the top 25 most abundant organism groups (overall, ~9%; Fig. 1, Table S1). When compared to baseline abundances in Lake Fargette, these results indicate enrichment of these groups, particularly CPR bacteria, which were barely detectable in the lake (Fig. 1b, Table S2).

**Fig. 1:**
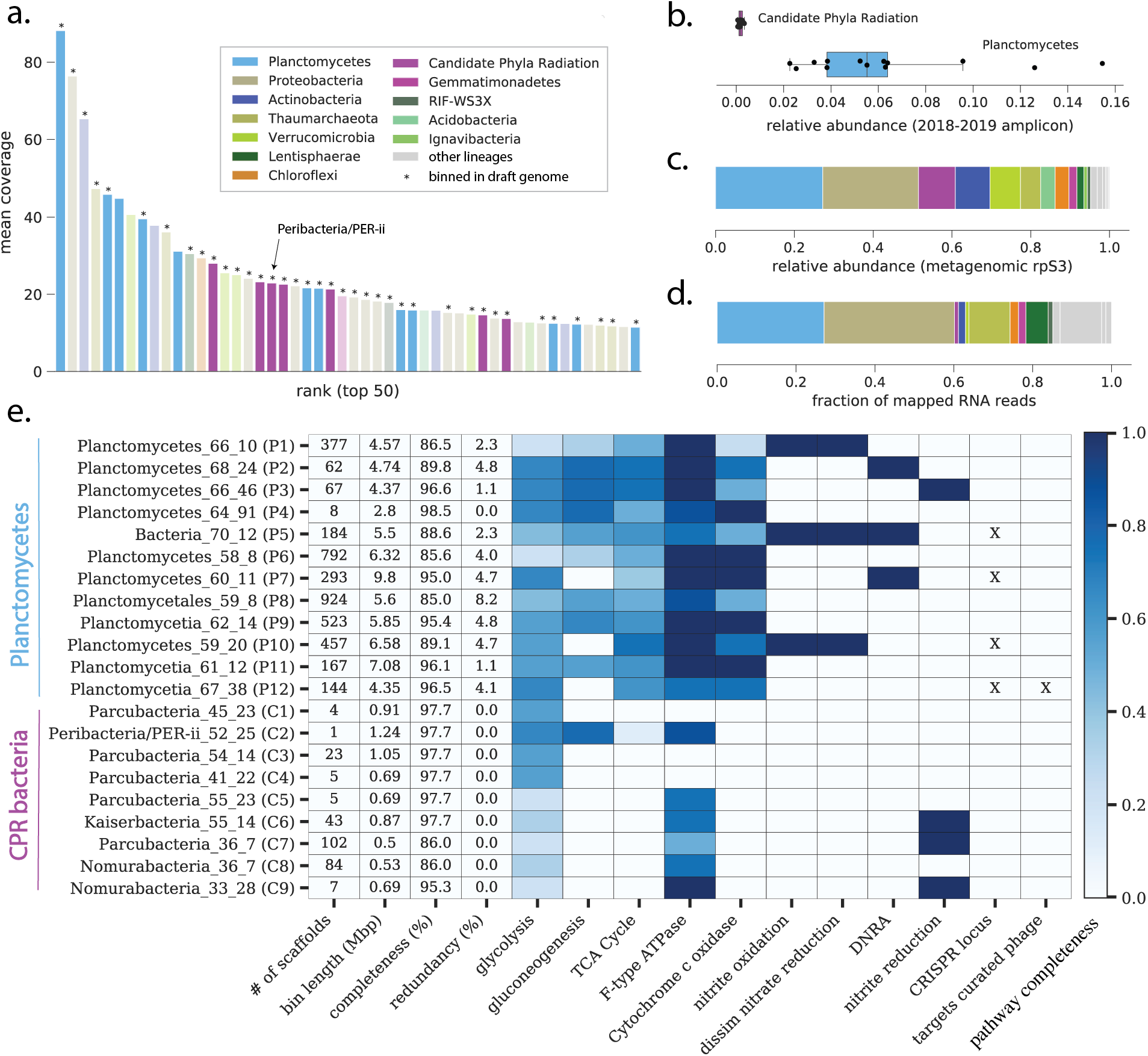
Long-term incubation enriched for members of the Planctomycetes and CPR bacteria. **a)** Rank abundance curve based on ribosomal protein S3 (rps3) coverage for organisms recovered at the end of incubation. Asterisks indicate marker genes that were binned into genomes. **b)** Relative abundance of CPR bacteria and Planctomycetes during monthly sampling of Lake Fargette. Each point represents the relative abundance of CPR bacteria or Planctomycetes in a given month. **c)** Overall community composition at the end of incubation, based on cumulative marker gene coverage. Panel **d** displays the fraction of RNA reads from the end of incubation that could be mapped to genomes. **e)** Sequence characteristics, metabolic predictions, and CRISPR-Cas loci information for genomes affiliated with the Planctomycetes and CPR bacteria. Cells with color fill indicate the fraction of key genes for each pathway (as defined by KEGGDecoder) that are present. “X”s indicates genomes with CRISPR loci, and if those loci contained spacers targeting at least one curated phage genome from the sample.

Where possible, scaffolds were assigned to genome bins that ranged in quality from draft to near-complete. The 48 genomes captured most of the phylogenetic diversity (38 of 50 most abundant rpS3 genes; Fig. 1a, Table S1/S3). Twelve genomes represent phylogenetically diverse Planctomycetes, including several from the large Planctomycetes and Phycisphaerae classes (Fig. S1, Table S3). Metabolic reconstructions suggested that the Planctomycetes are primarily heterotrophs with the potential to oxidize/reduce nitrite in some cases (Fig 1e). We also recovered 9 genomes of CPR bacteria, all but one of which came from lineages within the Parcubacteria (Fig. S1). The Parcubacteria genomes encoded minimal metabolic capacities consistent with symbiotic lifestyles [5]. However, several encoded a phylogenetically distinct *nirK* gene that may play a role in denitrification or energy conservation [6] (Fig. 1e).

One of the draft CPR genomes was for a member of the undersampled CPR lineage Peribacteria. We manually curated the original bin of 6 fragments into a single fragment that was circularized using a small, unbinned contig (Fig. S2, Materials and Methods). The newly reported 1,239,242 bp genome shares ~89% similarity in its 16S rRNA gene and is largely syntenous with a closely-related Peribacteria genome from Rifle, Colorado [7], supporting the accuracy of both assemblies (Fig. S3). Unlike the Rifle genomes, this Peribacter likely cannot synthesize purines *de novo*. Other notable differences include the presence of vacuolar-type H+/Na+-transporting ATPase complex and the lack of genes for biosynthesis of mevalonate. Based on biosynthetic deficiencies, we conclude that this bacterium probably relies on other organisms for many building blocks, but less so than the Parcubacteria. Supporting this idea, we observed that the Peribacter genome encodes all gluconeogenesis enzymes, including fructose-1,6-bisphosphatase I, which was not found in the Parcubacteria genomes (Fig. 1e).

### Phage and virus genomes

We reconstructed 12 phage sequences and 5 phage-like sequences, including three circularized genomes exceeding 100 kbp and two incomplete phage/phage-like fragments >300 kbp (Table S4). Phylogenetic analyses of encoded terminase and capsid proteins suggested that the phages likely fall within the Caudovirales, which is known to include numerous tailed phages with large capsids [8]. Additionally, we used phage gene content and analyses of bacterial CRISPR loci to infer that phages infect Planctomycetes, Proteobacteria, and Bacteroidetes (Fig. 2, Table S4, Materials and Methods).

**Fig. 2:**
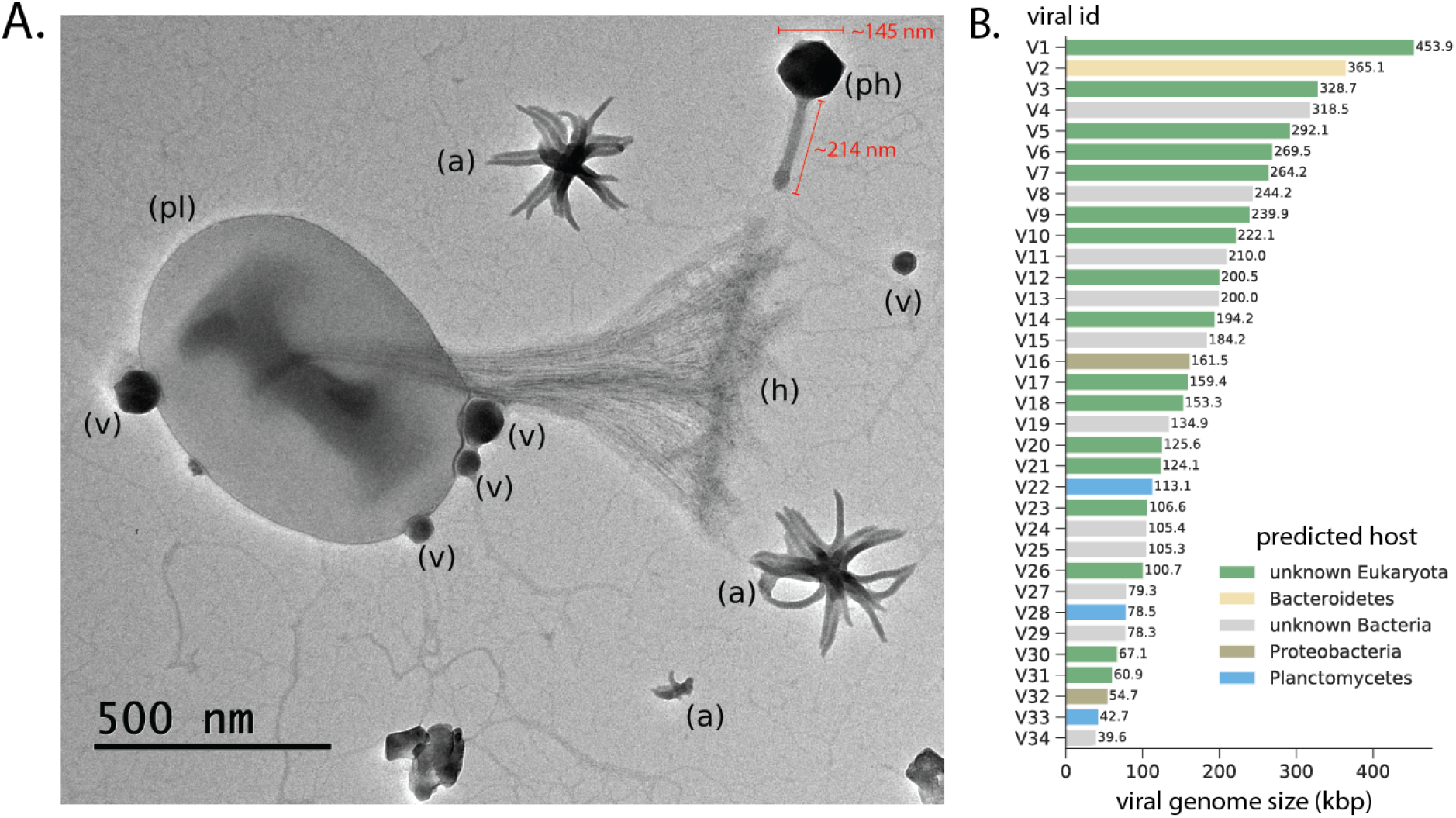
TEM imaging and viral genomics in the enrichment culture. **A.** Featured in this image is a cell inferred to be a Planctomycetes (pl) with a characteristic stalk and holdfast (h). Attached to the cell are four phage particles (v) (two different sizes, thus likely different phages). Also visible is one large tailed jumbo phage (ph) that is 145 nm in diameter with a 214 nm tail, as well as several aster-like nanoparticles (ALNs) (a). **B.** Genome sizes and predicted hosts for phages and eukaryotic viruses.

We also reconstructed large fragments of 17 eukaryotic viruses. Based on the phylogenetic placement of the major capsid protein and homologs of the Poxvirus Late Transcription Factor VLTF3, some viruses belonged to Iridoviridae (Betairidoviridae), Extended Mimiviridae, Phycodnaviridae, and Pitho-like viruses (Fig. S4-5, Table S4). Other viruses could only be classified at the superclade level, including those within a potentially novel Phycodnaviridae, Asfarviridae, Megavirales (PAM) clade, or did not contain either marker protein (Fig. S4-5).

### Imaging of incubation community

We imaged a diversity of cellular, cellular-like, and non-cellular particles using transmission electron microscopy (TEM) (Fig. 2, Fig. S6, Materials and Methods). Based on the presence of the extracellular ‘holdfast’ [9, 10], we infer that the cell imaged in Fig. 2 is likely a member of the Planctomycetes, with at least two attached distinct types of virus-like particles (VLPs). This finding is consistent with the high abundance of Planctomycetes in the enrichment as well as multiple phages predicted to infect them (~18-100x coverage).

We imaged numerous tailed phages (Fig. S6), including one with a 145 nm diameter capsid and 214 nm tail (“ph”, Fig. 2). This large capsid size is consistent with those of jumbo phages with genomes in the 300 kbp range [8], two of which were reconstructed here (Table S4). Despite these observations, TEM image counts suggested that the overall proportion of tailed phages decreased from 26% to less than 1 % of imaged objects over the course of the incubation (Table S5). Conversely, the proportion of ALNs increased substantially during the incubation, from 26% to 36% of imaged objects during the same period (Table S5).

### Transcriptomics of incubated community

Read mapping from a metatranscriptome collected with the metagenome suggested that Planctomycetes, and to a lesser extent, CPR bacteria, were active under the culture conditions, with Planctomycetes accounting for about 27% of RNA reads mapping to the non-redundant set of genomes (Fig 1d, Table S3). Interestingly, we detected very little transcription of genes from phage or eukaryotic viruses, suggesting that they were not actively replicating in the incubated community (Table S4). The lack of eukaryotes in the enrichment suggests that some of these particles may have derived from the inoculum and persisted for over two years.

## DISCUSSION

Planctomycetes are globally distributed across freshwater ecosystems where they are thought to play important roles in nitrogen and carbon cycling [11, 12]. Our analyses both expand genomic sampling for these organisms and suggest their interactions with other microbial groups, potentially including CPR bacteria. Although we cannot establish that the CPR bacteria [13] are symbionts of Planctomycetes, their co-enrichment suggests a possible association. Thus, the findings could guide future co-isolation of the CPR bacteria of the Parcubacteria and Peribacteria lineages, for which very little is currently known about host relationships [14].

It is also intriguing to find Planctomycetes and CPR bacteria co-enriched with ALNs. ALNs are enigmatic, organic femtoplankton entities that exhibit bloom-like behavior in various freshwater and coastal environments [3, 4]. Their defined morphologies suggest that they develop under genetic control; however, the genetic system responsible for producing ALNs remains unknown. Our co-enrichment data suggest ALNs may interact with or derive from co-occurring bacteria.

Overall, our results suggest that Planctomycetes might act as ‘hubs of interactions’ in subsystems within certain freshwater ecosystems.

## MATERIALS AND METHODS

### In vitro incubation, selection, and sequencing of microbial communities associated with ALNs

Lake Fargette is an artificial and highly eutrophic freshwater lake (surface area 1.2 ha, maximum depth 2.5 m) near Neuville in the French Massif Central (45°44’24”N; 3°27’39”E; 465 m altitude). Lake water was collected on March 15, 2017 when the highest density of ALNs was recorded in 2017 (9.0 ± 0.5 × 10^7^ ALNs·mL^-1^). Information about this sample is provided in Colombet et al. 2019 [3]. Within two hours after sampling, 100 L of raw lake water was filtered through a 25-μm-pore-size nylon mesh. Microbial communities were enriched by tangential-flow ultrafiltration using a Kross-Flow system (Spectrum, Breda, The Netherlands) equipped with a 0.2-μm cut-off cartridge. Aliquots (600 mL) of this concentrated 0.2 μm–25μm fraction were sequentially centrifuged at 8,000 g, 10,000 g (pellets discarded) then 12,000 g for 20 minutes each time at 14°C. Microbial communities contained in the final supernatant were cultivated for an initial period of 6 months at 4°C in the dark. The physico-chemical parameters of the starting sample are listed in Table S6.

After 6 months, the selected microbial community was enriched by centrifugation at 6000 g for 20 minutes at 14°C and the pellet (in liquid phase) incubated 25 more months at 4°C in the dark. At the end of the incubation, the medium was sonicated using Elmasonic S30 (Elma, Germany) and microbial communities were pelleted by centrifugation at 6000 g for 20 minutes at 14°C. The pellet suspended in distilled deionized water was used for nucleic acid extraction and amplification.

DNA and RNA extractions were performed using the RNA x DNA from soil kit (740143, Macherey-Nagel, Germany). The RNA sample was treated with Turbo DNA-free Kit (Invitrogen, Massachusetts, USA). The concentration of the samples was checked using a Qubit fluorometer (Invitrogen, Massachusetts, USA).

### TEM imaging of enrichment communities

Microbial communities were imaged at T0 and T31 to visualize their evolution, using a JEOL JEM 2100-Plus microscope (JEOL, Akishima, Tokyo, Japan) operating at 80 kV and 40,000x magnification. For each sample, 10 images were randomly captured in order to have a significant representation of microbial communities. We then defined the percentage of different observed phenotypes as a proportion of the total observed communities (Table S5).

### Monthly microbial community profiling at Lake Fargette

Samples were collected every month for 1 year (10/2018 to 09/2019) from the depth interval of 0-40 cm of lake Fargette. For each sample, microbial communities were collected on a 0.2 μm (Millipore) polycarbonate filter (until saturation, pressure < 25 kPa) and stored at −20°C until DNA extraction. The filters were covered with a lysing buffer (lysozyme 2 mg.ml-1 SDS 0.5%, Proteinase K 100 μg.mL-1 and RNase A 8.33 μg.mL-1 in TE buffer pH 8 at 37°C for 90 minutes. A CTAB 10% / NaCl 5M solution was added, and the samples were incubated at 65°C for 30 minutes. The nucleic acids were extracted with phenol-chloroform-isoamyl alcohol (25:24:1) the aqueous phase containing the nucleic acids was recovered and purified by adding chloroform-isoamyl alcohol (24:1). The nucleic acids were then precipitated with a mixture of glycogen 5mg/mL-1, sodium acetate 3M and ethanol 100% overnight at −20°C. The DNA pellet was rinsed with ethanol (70%), dried and dissolved in TE buffer. The DNA was purified with NucleoSpin^®^ gDNA Clean-up (Macherey-Nagel).

The amplification of the V4-V5 region of the bacterial small subunit rDNA was performed using the universal bacteria 515F and 928R primers modified with barcodes. PCR was performed in a total volume of 50 μL containing 1x final green reaction buffer, 2 mM final MgCl_2_, 0.2mM final dNTP, 100 μg.mL^-1^ final BSA,0.2 μM final of primers, 0.025 U.μL^-1^ final PROMEGA GoTaq HotsStart G2 and 5 μL of sample DNA.

To process the bacterial sequencing data (Illumina Miseq©, 2*250 bp), we used the FROGS pipeline [15]. After the clean-up procedure, sequences were assembled and clustered into OTUs with a similarity threshold of 95%. The representative sequences of each OTU were affiliated by similarity using the SILVA_132_16s database.

### Metagenomic/metatranscriptomic sequencing, binning, and marker gene analysis

Library preparation and sequencing of extracted DNA and RNA was performed at the Vincent J. Coates Genomics Sequencing Laboratory and Functional Genomics Laboratory at U.C. Berkeley. Extracted RNA was treated with the QiaSeq FastSelect kit to deplete rRNA. Samples were sequenced on a NovaSeq 6000 with 150 bp, paired-end reads. DNA and RNA were sequenced to a depth of ~10 Gbp and 25 million read pairs, respectively. Quality filtering and assembly of DNA reads, as well as annotation and binning of assembled scaffolds followed the workflow outlined in [16]. Genomes were classified at the phylum or class level based on the majority taxonomic affiliation of member contigs, and supplemented with BLAST searches of ribosomal proteins where necessary. Additionally, genome classification according to the GTDB scheme was assigned by running GTDB-tk with default parameters [17].

For the marker gene analysis (Fig. 1ac), we computed the mean coverage of scaffolds encoding a ribosomal protein S3 (rpS3) gene using bowtie2. Phylum-level taxonomic affiliation of these scaffolds was manually curated based on consensus gene-level taxonomy, binning information, and BLAST searches of protein sequence, where necessary.

### Quality filtering and genome curation for bacteria and archaea

Genomic bins were profiled for single copy genes using CheckM. For CPR bacteria, a custom workflow was used with a set of 43 marker genes sensitive to lineage-specific losses of ribosomal proteins in this group. We filtered all bins to those ≥ 70% completeness and ≤10% contamination and removed redundancy in the set at the species level using dRep (95% ANI) [18]. Retained genomic bins were manually ‘polished’ by removing contigs that were outliers in terms of GC content and coverage. Contigs with aberrant or ambiguous consensus taxonomy were manually expected if ≥ 5 kbp or automatically removed if <5 kbp.

For the Peribacteria genome (LakeFargette_0920_ALND_PER-ii_52_24), additional manual genome curation was performed. This involved fixing local scaffolding errors and extending the six original contigs using unplaced paired reads. The extended contigs were then joined based on end overlaps and paired read support. This resulted in two contigs, the larger of which was circularized and the other contained the 16S rRNA gene. Based on identification of a sequence within the larger contig that was also present at the end of the smaller contig, the larger contig was broken and joined to the smaller contig. This produced a single linear fragment. We used BLAST against the metagenome of the sample to identify a 4 kbp fragment that perfectly aligned to the ends of the linear fragment, circularizing the genome. The circularization overlap was trimmed and paired read support was verified by visualization throughout the entire final genome. Finally, the start position was adjusted based on the cumulative GC skew (Fig. S2). Genes/proteins were then re-predicted for the curated genome using Prodigal (single) [19].

We selected one of a set of near-identical complete Peribacteria genomes (SAMN03842449) from a previous publication [7] for comparison to the newly curated genome. Full genomes and 16S rRNA sequences were aligned in Geneious (Mauve plugin and MAFFT aligner, respectively).

### Phylogenomic and gene content analyses for CPR bacteria and Planctomycetes

For analyses of gene content, we focused on a subset of 21 genomes from the CPR bacteria and Planctomycetes. Gene/protein predictions were re-computed for this subset using Prodigal (single mode). Predicted protein sequences were annotated using kofamscan and resulting HMM hits were preliminarily filtered to those with e < 1×10^-6^ for use in downstream phylogenetic and metabolic analyses.

First, proteins annotated as rpS3 (K02982) were extracted and combined with reference sequences drawn either from NCBI (Planctomycetes) or a previous publication [20] on CPR phylogeny as well as a balanced sampling of other bacterial phyla as an outgroup. Each sequence set was aligned using mafft (*auto*) [21] and the resulting alignment was trimmed using trimal (*gt 0.1*) [22]. Maximum-likelihood trees were inferred using iqtree (*-m TEST -safe -st AA -bb 1500 -nt AUTO -ntmax 20*) [23] and subsequently visualized/annotated using iTOL [24].

For analyses of metabolism, we stringently filtered the kofamscan results, requiring that protein hits attain a score equal to or greater than the model specific thresholds for each KO. Results were then passed to KEGGDecoder [25] for visualization. For CRISPR-Cas analyses, we employed a custom script that first identifies repeat regions on scaffolds using PILER-CR [26] with default parameters, then detects Cas protein using the TIGRFAM HMM database [27] using hmmsearch [28] within 10 kbp (both upstream and downstream) of each identified repeat region. If at least one Cas protein was detected for a given repeat region, the spacer sequences from the adjacent CRISPR locus and also the paired reads and unplaced paired reads were extracted to identify spacer targets. Target identification was performed by blastn-short with an e-value threshold of 1×10^-3^. Hits were further filtered to those in which the spacer and targeted sequence were aligned over ≥90% of the spacer length and with no more than 2 mismatches.

### Viral genome identification and analysis

Viral genomes were identified through taxonomic profiling and identification of key viral structural proteins. While most recovered viral genomes were single-contig, in some cases multiple contigs of putative viral origin were binned together on the basis of GC content, coverage, and scaffold overlap. Viral genomes were tentatively classified as bacteriophage or eukaryotic viruses based on taxonomic profiling. Completeness and quality information for single-contig viruses was estimated using CheckV [29].

To establish phylogenetic affiliation of bacteriophages, we gathered curated sets of reference sequences for viral terminase and major capsid protein (MCP) from RefSeq. Reference sets were filtered to those sequences with sequence lengths 1±0.5 times the median and subsequently de-replicated at 95% identity using usearch (*-cluster_fast, -id 0.95*) [30]. Proteins for the newly identified phage genomes were predicted using Prodigal (meta mode). Terminase and MCP sequences were identified using BLAST (*-evalue 1e-20*) against a database built from the reference set. Sequences were concatenated with those of the reference set and aligned using MAFFT (*--reorder --auto*). Alignments were trimmed using trimal (*-gt 0.2*) and maximum likelihood trees were inferred using IQtree (*-m TEST -st AA*, version 1.6.12). Trees were decorated using taxonomic information for RefSeq sequences and putative lineages were assigned to newly identified phage genomes based on tree placement.

Host prediction for phages was performed using a combination of CRISPR-Cas targeting (described above) and taxonomic profiling. Briefly, for phylogenetic profiling, phage proteins were compared against UniRef100 using a custom DIAMOND database (*diamond blastp*) [31]. Hits were filtered to those with 70% or greater coverage of the query sequence and an e-value less than or equal to 1×10^-10^. We retrieved taxonomic affiliation for above-threshold hits and computed the percentage of genes on each phage with highest similarity to various bacterial lineages. If the bacterial lineage with the most hits among phage genes reached a percentage ? 3x that of the next highest lineage, it was assigned as the putative phage host. This method was previously shown to be consistent with host prediction via CRISPR-Cas targeting [32].

To infer the taxonomy of the 17 putative viruses of eukaryotes, all proteins were profiled against the Pfam database. 17 major capsid proteins (PF16903ļPF04451) from 11 eukaryotic virus bins and 8 VLTF3-like proteins (PF04947) from 8 bins served as markers. If there were multiple major capsid proteins per bin, the protein with the highest PFAM score was selected. The marker proteins were aligned with a reference dataset of capsid or VLTF3-like proteins from Nucleo-Cytoplasmic Large DNA Viruses [33] and 48 additional capsid proteins retrieved from NCBI (Extended Data File 1-2). The sequences were aligned, trimmed and the phylogenetic tree was constructed as described above. Finally, viruses were taxonomically classified based on phylogenetic placement.

### Transcriptome analysis

RNA reads were subjected to the same quality filtering pipeline referenced for DNA reads above and were subsequently mapped to the quality-filtered, de-replicated set of bacterial and archaeal genomes using bowtie2. To remove remnant rRNA and other non-mRNAs, we computed per-gene mapped read counts using pysam and removed those > 100 times the median non-zero per-gene read count for each genome. This process filtered about ~60 anomalously high-coverage ORFs, many of which corresponded to 16S rRNA or tRNA regions on scaffolds. Read counts were then aggregated by phylum-level lineage.

RNA reads were also mapped against the set of curated viral genomes described above. Mean coverage and coverage breadth were computed using CoverM (*genome mode, --min-read-percent-identity 0.95 --min-covered-fraction 0*).

### Data and software availability

Read data and draft genomes from this study are available through NCBI at PRJNA757735. Genome accession information for the 48 bacterial and archaeal genomes is also listed in Table S3. Custom code for the described analyses are also available on GitHub (github.com/alexanderjaffe/aln-enrichment). All supplementary figures, tables, extended data files, and genomes (including phage/viral genomes) are also available through Zenodo (https://doi.org/10.5281/zenodo.5362897).

## ACKNOWLEDGMENTS

We thank Alex Crits-Christoph, Adair Borges, Rohan Sachdeva, and Yue Clare Lou for informatics support and helpful discussions. We also thank Plateforme CYSTEM - UCA PARTNER (Clermont-Ferrand, France) for their technical support and expertise, the Innovative Genomics Institute and the Vincent J. Coates Genomics Sequencing Laboratory at UC Berkeley.

M.F., H.B., J.C., and T.S-N. performed sample collection, incubation experiments, and microscopy. A.L.J., L-X.C., M.C.S., and J.F.B. performed metagenomics and sequence analysis and J.F.B carried out the manual genome curation. All authors contributed to project design and manuscript writing.

